# From Likelihood to Fitness: Improving Variant Effect Prediction in Protein and Genome Language Models

**DOI:** 10.1101/2025.05.20.655154

**Authors:** Charles W. J. Pugh, Paulina G. Nuñez-Valencia, Mafalda Dias, Jonathan Frazer

## Abstract

Generative models trained on natural sequences are increasingly used to predict the effects of genetic variation, enabling progress in therapeutic design, disease risk prediction, and synthetic biology. In the zero-shot setting, variant impact is estimated by comparing the likelihoods of sequences, under the assumption that likelihood serves as a proxy for fitness. However, this assumption often breaks down in practice: sequence likelihood reflects not only evolutionary fitness constraints, but also phylogenetic structure and sampling biases, especially as model capacity increases. We introduce Likelihood-Fitness Bridging (LFB), a simple and general strategy that improves variant effect prediction by averaging model scores across sequences subject to similar selective pressures. Assuming an Ornstein-Uhlenbeck model of evolution, LFB can be viewed as a way to marginalize the effects of genetic drift, although its benefits appear to extend more broadly. LFB applies to existing protein and genomic language models without requiring retraining, and incurs only modest computational overhead. Evaluated on large-scale deep mutational scans and clinical benchmarks, LFB consistently improves predictive performance across model families and sizes. Notably, it reverses the performance plateau observed in larger protein language models, making the largest models the most accurate when combined with LFB. These results suggest that accounting for phylogenetic and sampling biases is essential to realizing the full potential of large sequence models in variant effect prediction.

## 1 Introduction

What do we learn when we fit a model to a distribution of natural protein or DNA sequences? A particularly important finding has been that the likelihood assigned to a sequence by such a model can serve as a zero-shot, nucleotide-resolution, measure of fitness. This insight is at the heart of methods which are transforming fields as diverse as therapeutic design, agriculture, materials science, pathogen forecasting and genetic diagnosis. And yet, at the same time, this finding is flawed.

A model trained on sequences from diverse organisms, whether whole genomes or protein sequences, learns patterns of nucleotide conservation shaped by natural selection. The likelihood thus provides a means of testing if a given sequence conforms to these patterns, and hence, whether or not it is likely to be functional. However, evolutionary forces alone do not fully determine the composition of the training distribution. It also reflects phylogenetic structure, historical contingency, and human biases in sequencing efforts. Given the success of using the likelihood as a measure of fitness, these issues have largely been treated as minor concerns, however recent advances in large-scale protein language models (pLMs) have brought them to the forefront. As pLMs have scaled to billions of parameters, they have achieved remarkable success in structure prediction and some generation tasks. Yet this scaling has not translated to improved performance in variant effect prediction. In fact, larger models appear to plateau or even regress in this task [Nijkamp et al., 2023, Gordon et al., 2024, Hou et al., 2025], revealing a widening gap between model likelihood and biological fitness (Fig. F.1).

There are at least three strategies to address the gap between likelihood and biological fitness: (1) modify the training data, (2) modify the model, and (3) modify the inference approach. In this work, we pursue the third strategy. We introduce the concept of likelihood-fitness bridging (LFB) and, using a simple model of selection and drift, propose a suite of fitness estimators that can be applied post hoc to any pre-trained protein or DNA language model. This approach offers a key practical advantage: it enables rapid exploration of new inference strategies without requiring the retraining of large models, making it highly efficient from a development standpoint. It is also computationally efficient at inference time – even simple LFB estimators, costing only ∼10× the runtime of a single forward pass, consistently outperform standard likelihood-based scoring. We assess how performance changes with scale for the ProGen2 and ESM-2 families of pLMs and find that LFB alleviates the previously reported performance plateaus, making the largest models the best-performing in both families. We also apply LFB to the Evo 2 whole-genome language models and observe consistent performance improvements, although without evidence of the same scaling trend.

In sum, we propose LFB as a general and computationally lightweight approach for improving zero-shot variant effect prediction.

## 2 Background

### 2.1 Protein and Genomic language models

Protein language models (pLMs) adapt methods originally developed for natural language processing to the domain of protein sequences. Proteins, represented as strings over a ∼20-letter alphabet corresponding to standard amino acids, provide a natural substrate for language modeling techniques. When trained on large databases of protein sequences sampled from across the tree of life, these models can uncover patterns of amino acid conservation shaped by millions of years of evolution [Meier et al., 2021, Brandes et al., 2023, Lin et al., 2023, Nijkamp et al., 2023]. This ability to model sequence constraints has enabled a broad range of downstream tasks: predicting 3D structure with high accuracy [Jumper et al., 2021, Baek et al., 2021], evaluating the effects of mutations [Meier et al., 2021], identifying functionally related proteins [Rives et al., 2021], and generating novel sequences with functional potential [Ferruz et al., 2022, Madani et al., 2023, Winnifrith et al., 2024]. As a result, pLMs (and other sequence models) are increasingly contributing to applied problems in disease risk prediction [Frazer et al., 2021, Gao et al., 2023, Cheng et al., 2023], drug discovery, and vaccine design [Youssef et al., 2025].

More recently, similar techniques have been extended to the modeling of entire genomes [Nguyen et al., 2023, 2024, Benegas et al., 2024, Dalla-Torre et al., 2025, Brixi et al., 2025]. Genomic language models (gLMs) aim to capture conserved patterns in DNA sequences, including non-coding regions. While still in the early stages of development, these models show great potential for tasks such as functional annotation, the design of regulatory elements, and whole-genome engineering [Consens et al., 2025, Benegas et al., 2025].

### 2.2 Likelihood based fitness estimation

To estimate the impact of a variant on protein function with a pLM or a gLM, it is standard to assume a monotonic relation between the probability of observing said variant and what is usually referred to as fitness. Concretely, take *p*_*θ*_ to be a model fit to a database of protein or DNA sequences. For predicting the effect of variants, it is usually assumed that the change in fitness, Δ*f*, can be estimated as,

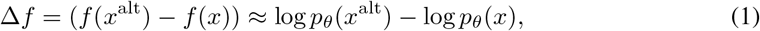

where *x*^alt^ is the variant sequence and *x* is the wild-type, reference, sequence [Hopf et al., 2017].

With masked language models such as the ESM family of pLMs [Meier et al., 2021, Brandes et al., 2022, 2023], computing the sequence log-likelihood, log *p*_*θ*_(*x*), is intractable. The standard approach is to use masked marginal predictions,

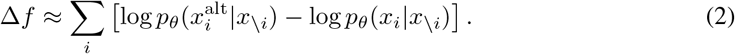

where *i* indexes over the positions in the sequence and *x* \ _*i*_ is the sequence *x* with the *i*-th amino acid, *x*_*i*_, set to the mask token.

In auto-regressive models such as the ProGen and ProtGPT pLMs [Nijkamp et al., 2023, Ferruz et al., 2022] and the Evo gLMs [Nguyen et al., 2024, Brixi et al., 2025], it is possible to obtain exact sequence likelihoods. The expression (1) can be computed by

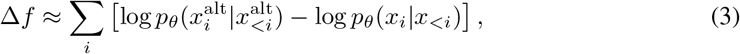

where *x*_*<i*_ is the sequence *x* up to index *i* − 1.

### 2.3 The gap between fitness and likelihood

Previous work has proposed a simple hypothesis for the relationship between the distribution of sequences and fitness, by describing evolutionary processes as a statistical physics system [Sella and Hirsh, 2005]. Within this formalism, a distribution of biological sequences can be described by a Boltzmann distribution *p*(*x*) ∝ *e*^*kf*(*x*)^. In this way log *p*(*x*) ∝ *f* (*x*) and fitness estimation based on likelihood is justified.

Recently, however, it has been recognized that there is a more complex relationship between fitness and likelihood. Biases in the composition of the training data affect predictions of fitness. Ding and Steinhardt [2024] show that pLM likelihoods are biased towards certain species due to acquisition bias in sequence databases. Gordon et al. [2024], Hou et al. [2025] observe that fitness predictions from pLMs suffer when the perplexity of the wild-type sequence under the model is too high or too low. The phylogenetic structure between extant sequences has also been shown to cause differences between likelihood and fitness even without the influence of sampling biases [Weinstein et al., 2022]. These have been proposed as reasons why larger protein language models, while fitting better to sequence databases, perform similarly or worse at zero-shot protein variant effect prediction than smaller counterparts [Weinstein et al., 2022, Nijkamp et al., 2023, Truong and Bepler, 2023, Bhatnagar et al., 2025] (Fig. F.1).

## 3 Methods

### 3.1 An alternative likelihood based fitness estimate

We propose a simple fitness estimate to overcome this gap based on existing model predictions using log-likelihood differences. We call this general strategy Likelihood-Fitness Bridging (LFB), Fig. 1.

**Figure 1.**
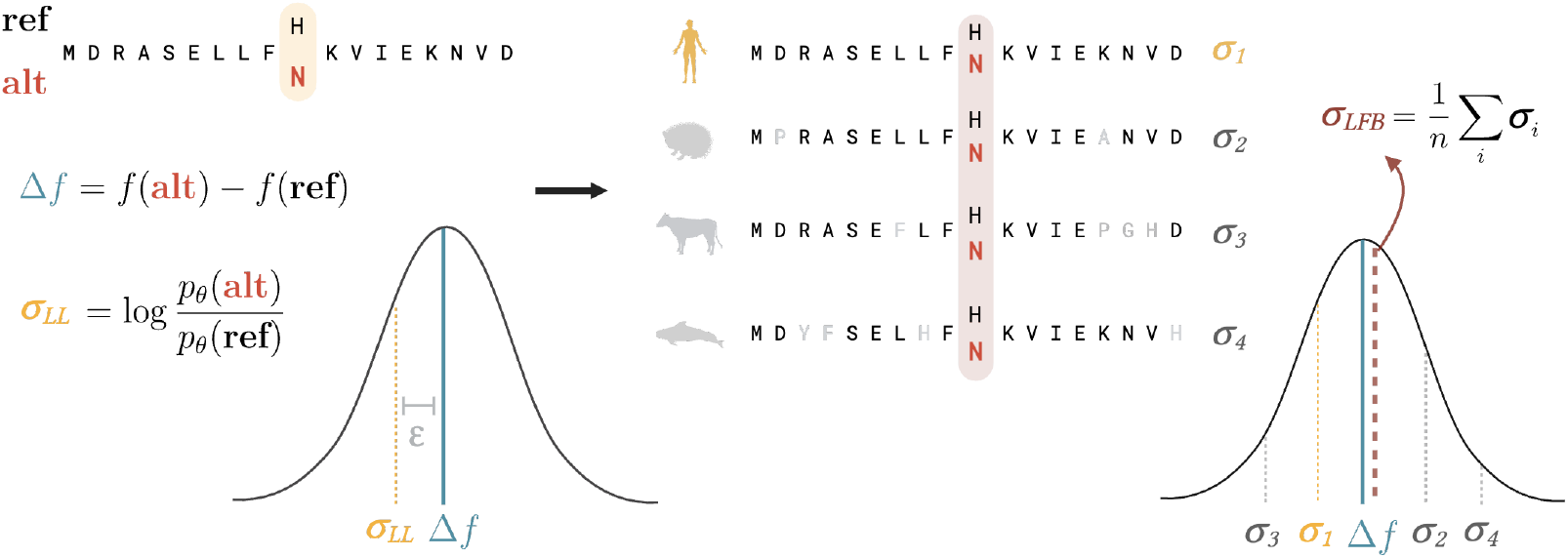
Overview of likelihood-fitness bridging (LFB) procedure. We propose that the log-likelihood ratio of alternative and reference sequences is a noisy estimate of the change in fitness Δ*f* (Left of arrow). By injecting the same substitution(s) into sequences that have resulted from similar evolutionary pressures, we can obtain multiple noisy estimates and hence their average, assuming independent noise, is a lower variance estimator of the true Δ*f* (Right of arrow).

The standard estimate of the impact of a variant *x*^alt^ of a sequence *x*, is

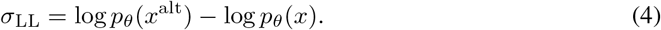

This single log-likelihood estimate *σ*_LL_ is noisy, at least affected by the phylogeny and composition of the training data. Therefore we instead “carry over” the reference and alternate alleles to related sequences, { *x*_*i*_ : *i* ∈ *I* }, which share a similar fitness landscape. Averaging the resulting differences in log-likelihood across these sequences reduces the effects of this noise,

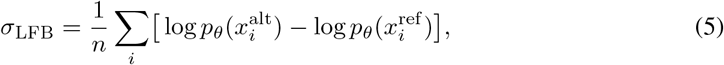

where 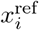 is the sequence *x*_*i*_ with the reference allele of *x* inserted (see Alg. 1).

The primary rationale behind this estimate is that closely related sequences, under similar selective pressures, will reflect a similar fitness landscape in their likelihoods.

If we regard the standard estimate *σ*_LL_ as a noisy estimate of the true change in fitness Δ*f* then, provided the fitness landscapes of the homologous sequences are suitably similar, the estimate *σ*_LFB_ should be unbiased, and provided the noise of each estimate, 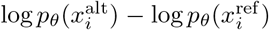, is independent, then the estimate *σ*_LFB_ should have lower variance. This method bears resemblance to test time augmentation approaches common in computer vision [Krizhevsky et al., 2012, Calvo-Zaragoza et al., 2020], but it differs in that it is part of an unsupervised method not attempting to get better likelihood estimates but departing from the likelihood in order to get better fitness estimates.

**A simple model of evolution** We study the behavior of our estimator under a simple model of molecular evolution including phylogenetic effects. As in Weinstein et al. [2022], we use the Ornstein– Uhlenbeck tree (OUT) process to model the evolutionary history of the present day sequences *x*_*i*_ used for LFB. We take *x*_*i*_ ∈ ℝ to be a continuous 1-d representation of these related sequences, all descended from some common ancestor according to a branching stochastic process.

We assume that across time, for this family of sequences, the fitness landscape has been constant and governed by 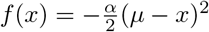, where *µ* is an optimal value for *x* and *α >* 0 determines the strength of selection, and that the sequences evolved according to

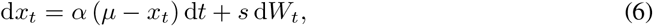

where *s >* 0 determines the strength of drift and *W*_*t*_ is a Wiener process [Butler and King, 2004]. If we assume that the common ancestral sequence follows the stationary distribution of this process, our present day sequences *x*_*i*_ can be expressed as

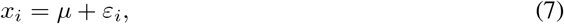

For 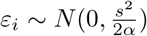. If we take *t* _*i,j*_ to be the time passed since the most recent common ancestor of *x*_*i*_ and *x*_*j*_, then we have

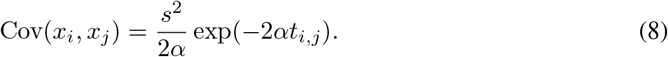

We hypothesize that models fit to databases of these observed natural sequences capture the contributions of drift, *ε*_*i*_, in their predictions, and that better fit models capture them more accurately. We formalize this by stating that around *x*_*i*_, the likelihood behaves such that log *p*_*θ*_(*x*) ∝ −(*µ* + *ε*_*i*_ − *x*)^2^, as opposed to matching the true fitness *f* (*x*) ∝ −(*µ* − *x*)^2^.

If we then consider the effect of a mutation – a perturbation 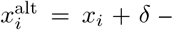 we can compute *E* [*σ*_LFB_] = *E [σ*_LL_] ∝ Δ*f* (see § E) meaning both *σ*_LL_ and *σ*_LFB_ are unbiased estimates of fitness up to scale by a constant. However, we also find that Var 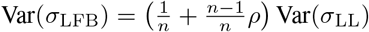, where *ρ* is the average correlation among the *x*_*i*_ (see §E).

Thus, under this model of selection and drift, our LFB fitness estimate has indeed lower variance than the standard estimate, as anticipated. Notably, this reduction is bounded by the average correlation among the sequences, so the reduction in variance provided by the LFB will be much greater when more phylogenetically disperse sequences are chosen for the averaging, which suggests a tradeoff between including close enough sequences which lie in the same fitness landscape and disperse enough sequences such that their errors are less correlated.

We hypothesize that sufficiently expressive models capture this phylogenetic structure of the data, represented above as noise^2^. One prediction that follows from this hypothesis is that larger models, with lower perplexities, should benefit more from LFB than smaller models in the same family.

### 3.2 Implementing our fitness estimator

For pLMs we found the sequences for the LFB procedure by making multiple sequence alignments with UniRef50 [Suzek et al., 2015] using MMseqs [Hauser et al., 2016]. By using the redundancy reduced UniRef50 we expect to find sequences which are different enough from each other to have less correlated predictions. We then filter these alignments in order to obtain homologous sequences which are suitably similar for LFB. We found a simple minimum percentage identity threshold of 30% performed best (Fig. F.2).

For the gLM, Evo 2, to obtain sequences for LFB we used the 447-way mammalian whole genome alignment from Zoonomia [Zoonomia Consortium, 2020]. We scored only coding-sequence variation, in order to compare with the pLMs. We randomly chose 9 species for each gene considered in addition to the human reference genome.

We outline the LFB algorithm in detail in (Alg. (1)). For the masked language models, ESM-2, we found that the unmasked-marginal scoring gave comparable performance to the standard masked-marginal scoring (Fig. F.3), so we used this more efficient method throughout unless specified. For ProGen2 and Evo 2 we used the standard log likelihoods. We include further implementation details in (§D).

## 4 Results

### 4.1 Baselines

Our goal is to assess whether augmenting a generative model with likelihood-fitness bridging (LFB) improves the ability to predict the impact of genetic variation on fitness. We do so using two classes of tasks – classification of variants with known “benign” and “pathogenic” clinical labels, and correlation with deep mutational scanning (DMS) measurements from a large number of experiments designed to measure fitness (or closely related properties). For both tasks, we use publicly available curated data from ProteinGym [Notin et al., 2022, 2023].

To establish families of generative models as baselines, we first consider families of pure sequence pLMs (*i*.*e*. we don”t consider hybrids such as those which also leverage 3D structure information) for which at least two different model sizes have been made publicly available. This gives us five families; CARP [Yang et al., 2024], ESM-2 [Lin et al., 2023], ProGen2 [Nijkamp et al., 2023], RITA [Hesslow et al., 2022] and Tranception [Notin et al., 2022]. Fig. F.1 compares the performance of these models as measured by the weighted average^3^ spearman across all DMS. Consistent with previous reports, we find the largest models exhibit a plateau in performance. Since the ESM-2 and ProGen2 families contain both the largest models and also the best performing models, we take these families as our baselines and also use them as case studies for exploring the benefits of likelihood-fitness bridging. An added benefit is that these two families are complementary, differing in design and training in a number of important ways. For instance the ESM-2 family is trained for masked-language modeling, while ProGen2 uses an auto-regressive decoder. Analyzing the impact of likelihood-fitness bridging in the context of both families in parallel therefore enables us to explore the sensitivity of the approach to the underlying model.

Finally, nothing about our approach is specific to pLMs and so we explore the benefits of LFB for gLMs as well. To do so we use the Evo 2 family Brixi et al. [2025].

### 4.2 Predicting Disease-Causing Variants

Our first task is to assess if a model can separate variants which have been seen in the human population and classified as “benign”, from those thought to significantly increase the risk of disease, “pathogenic”. As recommended in [Dias et al., 2024] we assess model performance by computing the area under the receiver-operating characteristic curve on a gene-by-gene basis and then compute the average across genes. We restrict our attention to genes for which there are at least 10 Benign and 10 Pathogenic labels, giving us a total of 305 disease-associated genes. For both the ESM-2 and ProGen2 families, the smallest models, when combined with LFB outperform larger models using the log-likelihood (Fig. 2a, Fig. F.4a, Table 1). We also see that although all models perform well when combined with LFB the largest of the pLM models is the best performing. The Evo 2 gLMs also benefit from LFB, although the performance gain is more modest and the scaling trend is not reversed. A possible explanation for this contrast in behavior to pLMs is that the primary performance limitations of this family do not arise from the phylogenetic structure of the training data. A full picture of performance gains for all models is shown in Fig. F.5.

**Table 1.** Performance of different models on DMS and Clinical Labels. ↑ indicates higher is better.

| Model Class | Size | Mean Spearman on DMS ↑ | | Mean AUC on Clinical Labels ↑ | |
| --- | --- | --- | --- | --- | --- |
|  |  | baseline | LFB | baseline | LFB |
| ESM-2 | 8M | 0.250 | 0.355 | 0.653 | 0.889 |
|  | 35M | 0.325 | 0.388 | 0.751 | 0.904 |
|  | 150M | 0.395 | 0.416 | 0.839 | 0.920 |
|  | 650M | 0.417 | 0.428 | 0.895 | 0.933 |
|  | 3B | 0.404 | 0.430 | 0.903 | **0.938** |
|  | 15B | 0.398 | **0.436** | 0.895 | **0.938** |
| ProGen2 | 151M | 0.339 | 0.361 | 0.873 | 0.888 |
|  | 764M | 0.379 | 0.404 | 0.894 | 0.923 |
|  | 2.7B | 0.380 | 0.403 | 0.887 | 0.920 |
|  | 6.4B | 0.389 | 0.433 | 0.869 | 0.930 |
| Evo 2 | 7B | - | - | 0.837 | 0.850 |
| Evo 2 base | 1B | - | - | 0.868 | 0.878 |
|  | 7B | - | - | 0.845 | 0.856 |
|  | 40B | - | - | 0.839 | 0.853 |

**Figure 2.**
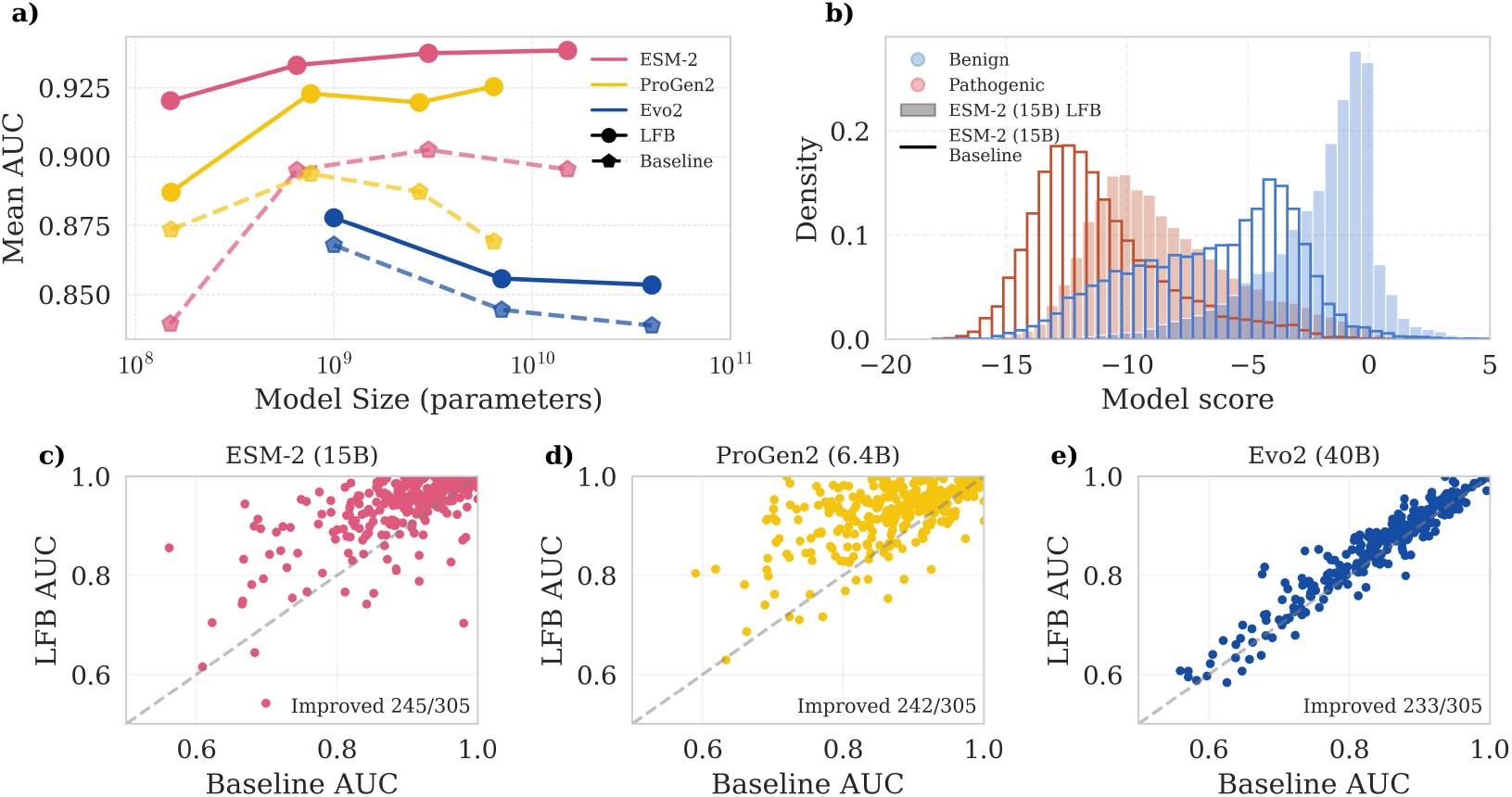
Comparison of pLM and gLM families with and without likelihood-fitness bridging at clinical label prediction. a) Average AUC comparison of models in ESM-2, ProGen2 and Evo 2 (base) families, with and without LFB (see Fig. F.4a for bootstrap error bars). b) Distribution of scores of all variants in the assessment for ESM-2 15B with and without LFB (see Fig. F.6 for all other model sizes and families). c), d), e) Performance comparison of ESM-2 15B (c), ProGen2 XL (d) and Evo 2 7B (base) (e) with and without LFB on a per-gene basis.

When comparing the score distributions of all clinical variants between ESM-2 15B and its LFB-augmented counterpart (2b, see also Fig. F.6), we observe that the LFB fitness scores achieve a better separation between “benign” and “pathogenic” labels with the benign variant distribution exhibiting a reduced low-score tail. The LFB procedure also shifts the overall distribution to more positive scores, although this appears to be a result of the unmasked-marginal scoring, under which the scores are generally more negative than under masked-marginal scoring (Fig. F.6).

When comparing performance on a gene-by-gene basis for the largest models in the ESM-2 and ProGen2 families, we see that almost perfect separation of “benign” and “pathogenic” labels is achieved for many genes (Figs. 2c and 2d). This will impact how the model may be used in the clinical setting. Clinical variant annotation proceeds by combining multiple sources of evidence, such as population frequency, family incidence, or scores from variant effect prediction models [Richards et al., 2015]. Each source of evidence is weighted according to the quality of the evidence, and guidelines have recently changed for variant effect predictors by accounting for their performance [Pejaver et al., 2022]. We can anticipate that given LFB near-perfect performance for a number of disease-associated genes, this approach will be valuable for clinical annotation.

### 4.3 Assessing Concordance with Experimental Assays

Complementary to clinical variant prediction, another approach to assess fitness prediction performance is by comparison with deep mutational scans (DMS). ProteinGym consists of manually curated DMS assays, spanning 186 proteins and measuring the impact of ∼2.5M variants on protein function with a range of assay types. In many cases, however, the assay provides an incomplete picture of whether or not the protein is functioning properly. For instance, an assay might measure protein stability but not binding affinity. In addition, some experiments have a modest correlation between replicates. Thus, even a perfect fitness prediction model will not exhibit a perfect correlation with functional assays. Nevertheless, by considering a large number of assays and proteins, we expect that the average performance across these assays should be a reasonable means of assessing if one model is a better predictor of protein fitness than another. In practice, to compute the mean spearman values across all models, we subsampled 200 measurements from each DMS.

A comparison of model performance of the ESM-2 and ProGen2 families with and without LFB is shown in Fig. 3 (Table 1, Fig. F.4b, Fig. F.7 and Fig. F.8). We did not compare the performance with Evo 2, as predictions are unavailable for many assays. LFB results in performance gains for ESM-2 models – the 8M parameter model with LFB outperforms the original 35M parameter, the 35M model with LFB matches the original 150M model and the 150M model with LFB matches the original 650M model. Notably, while further scaling of the original ESM-2 models resulted in decreasing performance, the likelihood-fitness bridged 8B and 15B parameter models continue to improve, with the largest model now also being the best performing. Similarly the ProGen2 Medium size model with LFB outperforms the original XL model and the XL model with LFB is the best-performing model in the family (Table 1).

**Figure 3.**
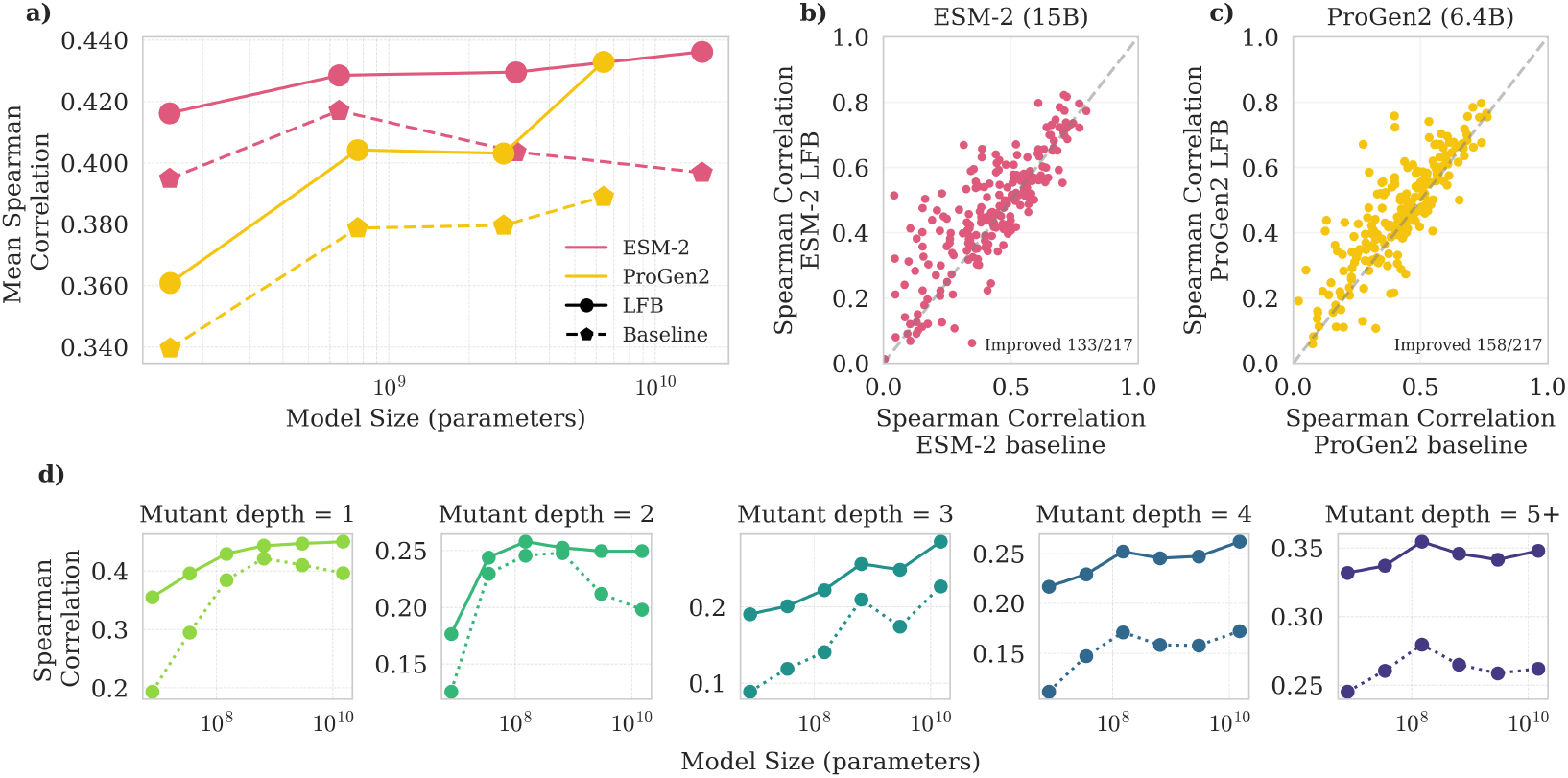
Comparison of pLM families with and without likelihood-fitness bridging at fitness estimation as measured by correlation with DMS. a) Comparison of all models in ESM-2 and ProGen2 families, with and without LFB (see Fig. F.4b for bootstrap error bars). b) Comparison of ESM-2 15B model with and without LFB on a per-experiment basis. c) Comparison of ProGen2 XL model with and without LFB on a per-experiment basis. d) Mean Spearman correlation on variants at different mutation depths for the ESM-2 family of models. In a), b) and c), correlations are taken across all 217 DMS, randomly subsampling to at most 200 variants per assay, and in a) the mean is weighted as in Notin et al. [2023].

Comparing the performance with and without LFB on a per-experiment basis (Fig. 3b and c), we see that the average performance boost observed in Fig. 3a for both the ESM-2 and ProGen2 models comes from broad performance gains across most experiments (see also Fig. F.7). These gains span assay type (Activity, Binding, Expression, Organismal Fitness) (Fig. F.9), alignment depths (Fig.F.10), and diverse DMS-types and proteins more generally (Fig. F.8).

Critical to understanding the potential impact of LFB on design tasks, we explore its impact at varying edit distances, by focusing on measurements probing combinations of mutants. Although scaling trends are less clear for multiple mutants, LFB consistently improves performance across all mutation depths (Fig. 4d).

**Figure 4.**
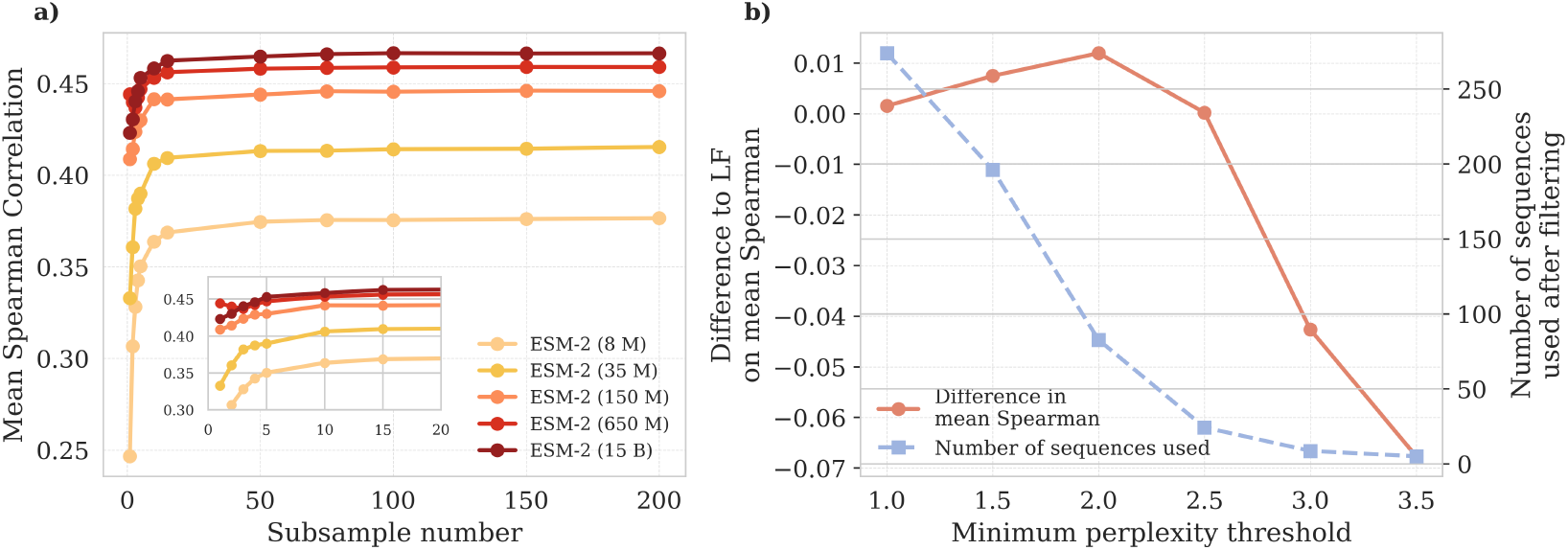
Scalability and relation to perplexity. a) Mean Spearman values for LFB estimators, using random subsamples of different sizes of the sequences used for the averaging. b) Comparison between the standard ESM-2 15B LFB model, and an LFB estimator obtained by further filtering the sequences used by their minimum perplexity, and with a maximum perplexity of 10. On the y axis, the difference in mean Spearman across DMS and also the number of sequences after filtering by both sequence identity and perplexity. Correlations are taken across all 217 DMS without subsampling variants, and the mean is unweighted.

### 4.4 Efficiency of the method

In order to better understand the compute-performance tradeoff of LFB we took random subsamples of decreasing sizes of the sequences used in the averaging procedure and produced LFB estimates with these reduced alignments (Fig. 4a). We found across the ESM-2 family that we retain most of the benefit of LFB with as few as 10 sequences. Given that we only require one forward pass per sequence with the ESM-2 model using unmasked-marginal scoring (Fig. F.3), this provides an extremely scalable variant effect prediction method.

### 4.5 Relationship to perplexity

Recent works have identified a trend between the perplexity of a sequence under a generative model and the performance in estimating fitness [Gordon et al., 2024, Hou et al., 2025]. For the top-performing model on the DMS benchmark, we tested whether filtering the sequences used for LFB by estimates of their pseudo-perplexities could improve the resulting fitness estimate (Fig. 4b, Fig. F.11). We used the single forward pass pseudo perplexity calculation developed in Gordon et al. [2024]. We find that filtering for sequences with a perplexity of at least 2 provides further improvement to the performance. However, filtering to even larger perplexity values results in a decline in performance, suggesting a tradeoff between model confidence and lack of information, echoing findings by [Gordon et al., 2024] and [Hou et al., 2025].

## 5 Discussion

Recent works have suggested that in order to bridge the gap between likelihood and fitness we should place greater emphasis on the distribution of sequences provided to the model during training [Nijkamp et al., 2023, Ding and Steinhardt, 2024, Gordon et al., 2024], however this approach has limitations. First, the relationship between model design and optimal data for training is poorly understood, making the process of data selection challenging. Second, the optimal choice of data selection will depend on the downstream task. Hence, models will need to be trained with downstream tasks defined from the outset, thereby limiting their potential in both multi-task learning and domain adaptation. Another approach is to modify the model, such as by joint modelling of fitness and phylogeny. However as discussed in Weinstein et al. [2022], there is a non-identifiably issue.

In this work we explore a third strategy. Rather than changing the underlying model”s training data, or modifying the model building approach, we instead propose that models be built to extract fitness predictions from a preexisting pLM of gLM. This is similar in spirit to Low-Rank Adaptation (LoRA) style fine-tuning [Hu et al., 2022] or some retrieval mechanisms (such as in Tranception [Notin et al., 2022]), where the language model remains unaltered (and hence its potential to perform diverse tasks unhindered) but is instead augmented to perform a specific task.

We find that the largest pLMs benefit the most from LFB, which is consistent with the idea that they are achieving lower perplexities but worse fitness prediction by learning both fitness constraints and phylogenetic relationships. In contrast, while our exploration of gLMs is limited to a subset of the Evo 2 family, it appears that at least for these cases, capturing phylogeny is not the primary cause of the gap between likelihood and fitness. Instead we see comparable performance gains for both models with LFB and the smaller model continues to be stronger. So while LFB improves performance, in this case the relationship with phylogeny is less clear.

### Limitations

The LFB estimators proposed in this work are intentionally simple and serve as a starting point for more sophisticated inference strategies. While motivated by a model of selection and drift, the current implementation does not explicitly incorporate the underlying phylogeny of the sequences used for LFB. Another important limitation is the focus on single or combinatorial substitutions; insertions and deletions (indels) are not included in the current framework. Furthermore, while multiple details of the implementation likely reduce the impact of sampling biases, these biases are not explicitly modelled. Finally, while LFB performs well across a wide range of benchmarks, its performance has so far only been validated on coding regions. Extensions to non-coding regions remains to be explored.

## 6 Conclusions

The surprisingly close connection between fitness and the distribution of natural sequences has enabled powerful zero-shot variant effect prediction by simply using likelihoods from generative sequence models to estimate fitness. However with sufficiently expressive models it seems we are reaching the limitations of this connection, and as our ability to model the distribution of protein sequences improves, a gap between likelihood and fitness is becoming apparent. In this work we propose a framework for improving variant effect prediction with protein language models by bridging this gap. We adapt the theory developed in Weinstein et al. [2022] and use it to describe the evolutionary history of sequences in a neighborhood of interest. Under such a model of evolution, sufficiently expressive models will be able to capture the effects of genetic drift, hence their likelihood will be a suboptimal fitness estimator, and according to this theory, LFB should provide a better estimate. When LFB is applied to the ESM-2, ProGen2 and Evo 2 families, all models enjoy performance gains. Furthermore, in both the ESM-2 and ProGen2 families, the largest model performs best once combined with our bridging model. This is consistent with the idea that the largest models are starting to capture non-fitness related structure in the data and suggests that further scaling of these models will result in additional performance gains when combined with LFB.

The performance gains span the majority of tested proteins and also span assays probing different aspects of fitness, suggesting that the benefits of bridging will apply to a broad range of downstream applications. We found LFB to improve fitness prediction at multiple edit distances, suggesting its potential for design tasks. And from a clinical impact perspective, we observe broad and often large improvements in performance. We are therefore optimistic that the use of likelihood-fitness bridging will result in better understanding of the genetics of disease, improve preventative care and increase the diagnostic yield of patient sequencing.

This work supports the hypothesis that variant effect prediction can be improved by taking into consideration the fact that natural sequence distributions are most likely the result of a combination of fitness, phylogeny and various sampling biases. While there are many promising directions for incorporating these factors, LFB has the advantage of applying to preexisting models without requiring retraining, making it a practical and scalable addition to current inference workflows. The approach proposed here is simple but also neglects a number of important considerations. Thus, we see this work as a promising starting point for a richer class of inference strategies that reconcile evolutionary modelling and modern sequence-based machine learning.

## Acknowledgments

We would like to acknowledge the reviewers whose thoughtful comments helped improve the work. We thank the Scientific IT team at the Centre for Genomic Regulation (CRG) for their assistance with computational infrastructure, in particular Emyr James and Emilio Palumbo. We are also grateful to the CRG Core Technologies Programme for their assistance. We thank other members of the Dias and Frazer lab for their thoughtful feedback and many interesting discussions throughout the development of this work. PN is supported by a fellowship from the “la Caixa” Foundation ID 12070017, as part of the funding received from the European Union”s Horizon 2020 research and innovation programme under the Marie Skłodowska-Curie grant agreement No. 713673. This work was supported by the Spanish Ministry of Science and Innovation (PID2022-140793NA-I00 and PID2022-143210NA-I00 both funded by MCIN /AEI /10.13039/501100011033 /FEDER, UE). We acknowledge support of the Spanish Ministry of Science and Innovation through the Centro de Excelencia Severo Ochoa (CEX2020-001049-S, MCIN/AEI /10.13039/501100011033), and the Generalitat de Catalunya through the CERCA programme.

## A Code Availability

The code to run LFB is available at https://github.com/DiasFrazerGroup/lfb. ESM-2 models are available from https://github.com/facebookresearch/esm, ProGen2 models at https://github.com/enijkamp/progen2 and Evo 2 models at https://github.com/ArcInstitute/evo2. The ProteinGym code and data can be found at https://github.com/OATML-Markslab/ProteinGym.

## B Computational resources

We ran all models in an HPC setting. We used 1 Nvidia H100 GPU for gLM and pLM inference. Memory requirements didn”t exceed 35GB RAM. In order to process alignments with MMseqs we ran jobs in parallel with 10 CPU cores, and 35GB RAM.

## C Impact Statement

This paper introduces a framework for enhancing the performance of large language models at predicting the effect of variants on protein and DNA function and human health. By advancing variant effect prediction, this work has the potential to drive progress across multiple fields, from protein design for therapeutics and bioengineering, to clinical genetics. While the insights of this model can guide diagnostic care and help uncover the genetic architecture of disease, its predictions should complement – not replace – experimental validation and expert interpretation. In this sense, ethical considerations include transparency and interpretability to ensure the responsible usage of the model in real life applications. One of the benefits of approaches that train on the whole protein or DNA universe, is the robustness to biases in human genetic studies, and therefore better generalization across genetic ancestries. Nevertheless, as with any AI-driven approach, care must be taken to ensure equitable benefits across populations and to prevent misuse in genetic profiling. Nonetheless, this work primarily seeks to enhance computational methods for studying protein and DNA variant effects, with no foreseeable direct societal harm.

## D D Implementation details of LFB

### D.1 Alignments

To produce protein sequence alignments we use the MMseqs search tool [Hauser et al., 2016] against the UniRef50 database [Suzek et al., 2015]. We used the arguments: -s 7.5 –num-iterations 5.

To produce the DNA sequence alignments for the human clinically annotated variants, we used the unprocessed DNA level variants provided in ProteinGym. To obtain sequences from other species we used the Zoonomia 447-way primate and mammalian alignment. We used HAL liftover to map the variants from the human reference genome to these genomes [Hickey et al., 2013]. Then we extracted 8,192 length segments centered around the variant at these genomes, to obtain the same context length around variants as in Brixi et al. [2025].

### D.2 Log-likelihoods from pLMs

For ESM-2, in place of (2) we use”

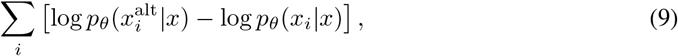

where *i* indexes over amino acid position in a protein sequence *x*. This scoring system has been shown to perform similarly to other masked language model scoring systems in Meier et al. [2021], and Gordon et al. [2024] outline reasons why BERT trained models still make predictions when conditioned on a fully unmasked sequence. We found it performed similarly in practice to the masked-marginal scoring (Fig. F.3), and it only requires one forward pass for each sequence.

For ProGen2, and for Evo 2 we use the log-likelihoods as in eq. (3), but also ensemble over the sequences in different directions. For ProGen2 we average over the prediction for the sequence and the reversed sequence,

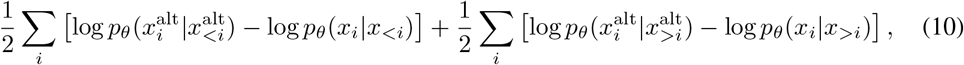

which is possible as the model is trained also on reversed sequences. For Evo 2 we average over predictions for the sequence and its reverse complement *y*,

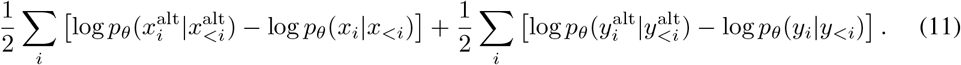

### D.3 LFB algorithm

We describe in the below algorithm how to produce a LFB estimate given an alignment of a reference sequence against related sequences, and generative model capable of scoring these sequences.

#### Algorithm 1

Scoring variants with LFB

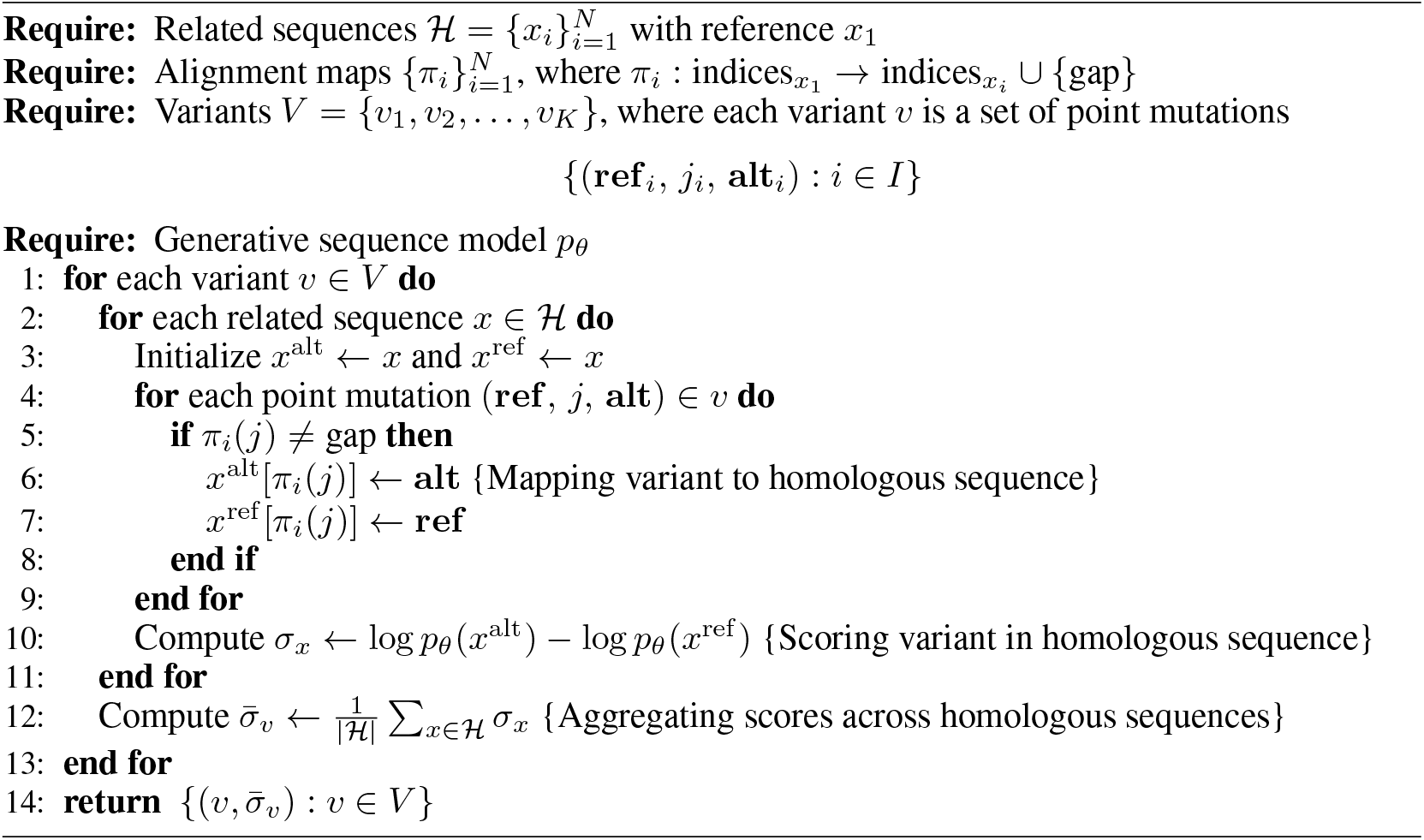

Notably, we only need the alignment mapping on those positions of the reference and alternative alleles. One consequence of this algorithm is that if no variants are mapped over (due to gappy alignment, or the variants being in excluded domains or less important regions), the difference in log-likelihood will vanish. We tried also averaging only over non-gap sequences for each position, but found this had a slightly negative impact (Fig. F.12). Another notable choice is the inclusion of sequences in the average with wild-type alleles different to the reference sequence. We tried only averaging over those sequences which matched the reference allele for each position, and similarly found slightly diminished performance (Fig. F.12).

## E Sketch proof of lower variance under OUT model

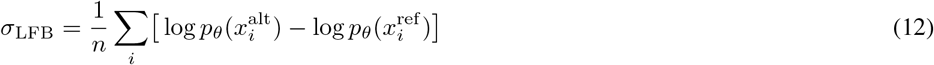

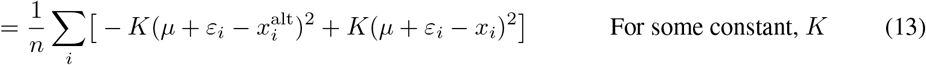

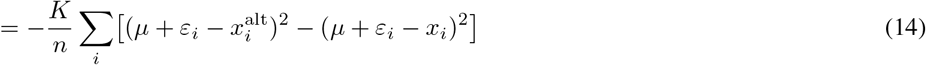

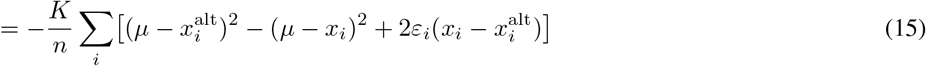

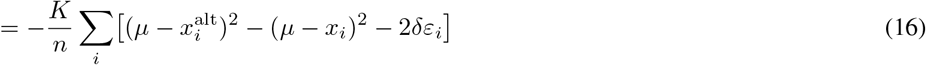

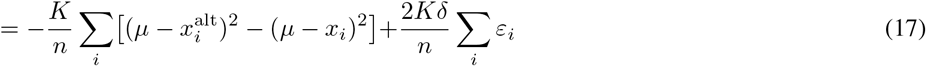

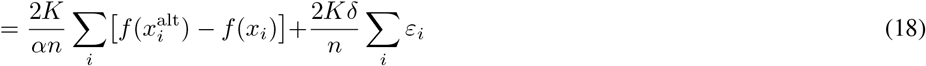

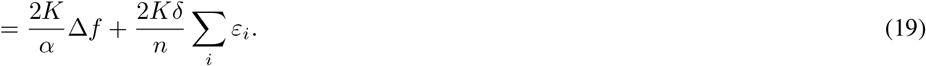

Whereas, for the single log-likelihood calculation we have

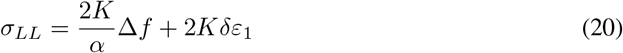

Therefore, we find

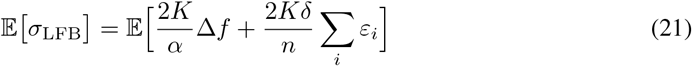

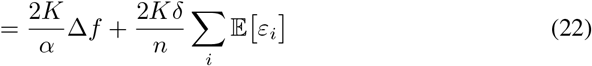

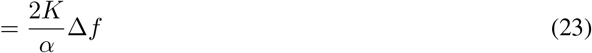

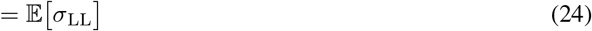

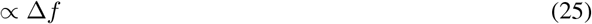

Similarly,

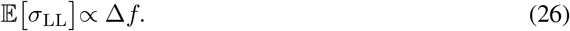

And considering the variance we have

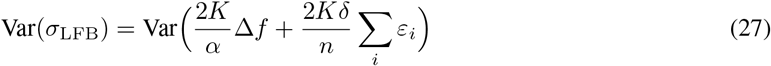

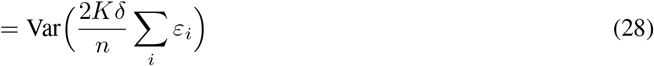

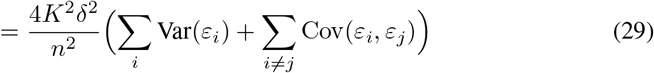

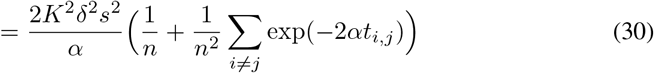

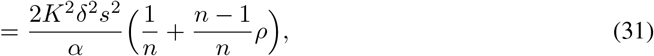

Where

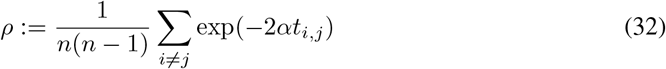

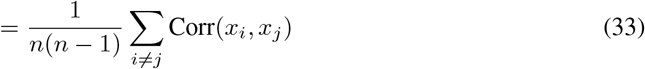

the average correlation among the *x*_*i*_, or equivalently the *ε*_*i*_.

And similarly we have

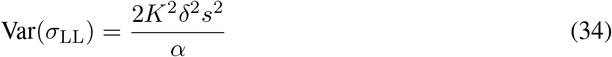

So

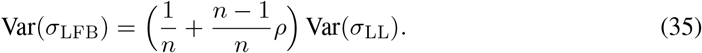

## F Supplementary figures

**Figure F.1.**
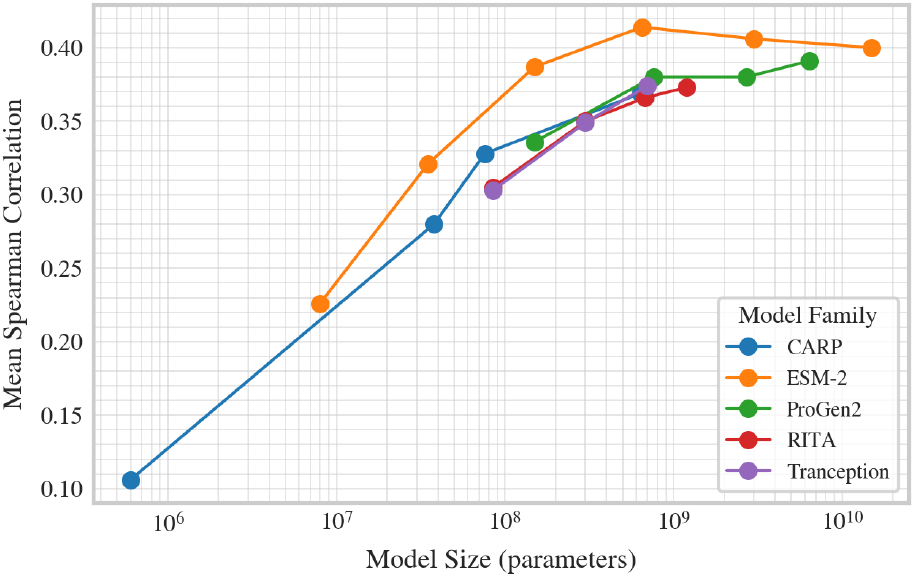
Fitness estimation scaling of candidate baseline families. Performance assessment of five protein language model families at variant effect prediction, as measured by mean correlation with deep mutational scanning assays, plotted against the number of parameters of each model. For smaller models, increasing model size results in better performance but for larger models, the performance plateaus, or decreases. Results were taken from https://proteingym.org/benchmarks.

**Figure F.2.**
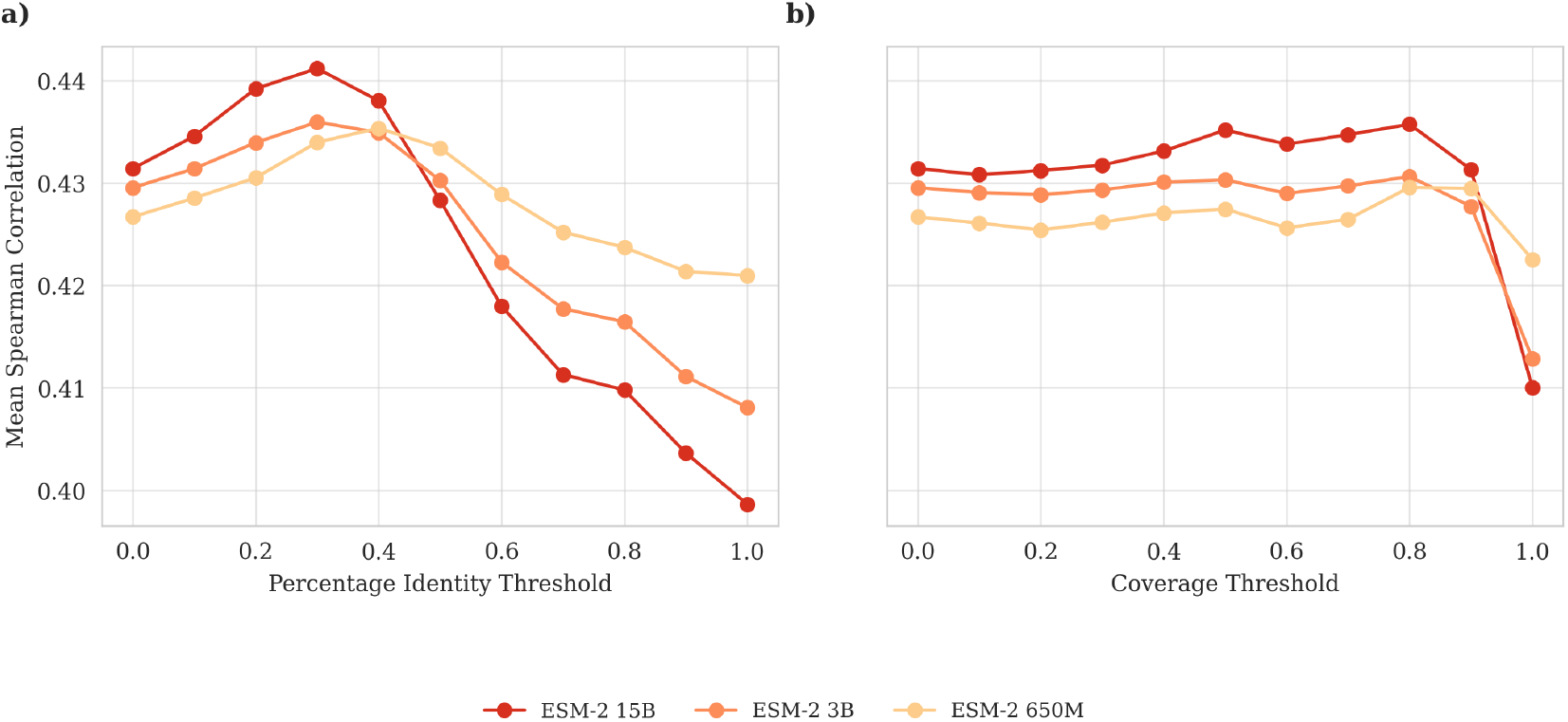
LFB performance across different alignment filtering strategies, measured by mean spearman correlation to DMS experiments. (a) Mean Spearman correlation as a function of minimum percentage identity threshold in MSA filtering. (b) Mean Spearman correlation as a function of minimum coverage threshold in MSA filtering. Each line represents a different ESM-2 model: 15B (dark orange), 3B (medium orange), and 650M (light orange). Unmasked-marginal scoring is used and mean correlations are taken across all the 217 DMS without subsampling of the variants, and the mean is weighted as in Notin et al. [2023].

**Figure F.3.**
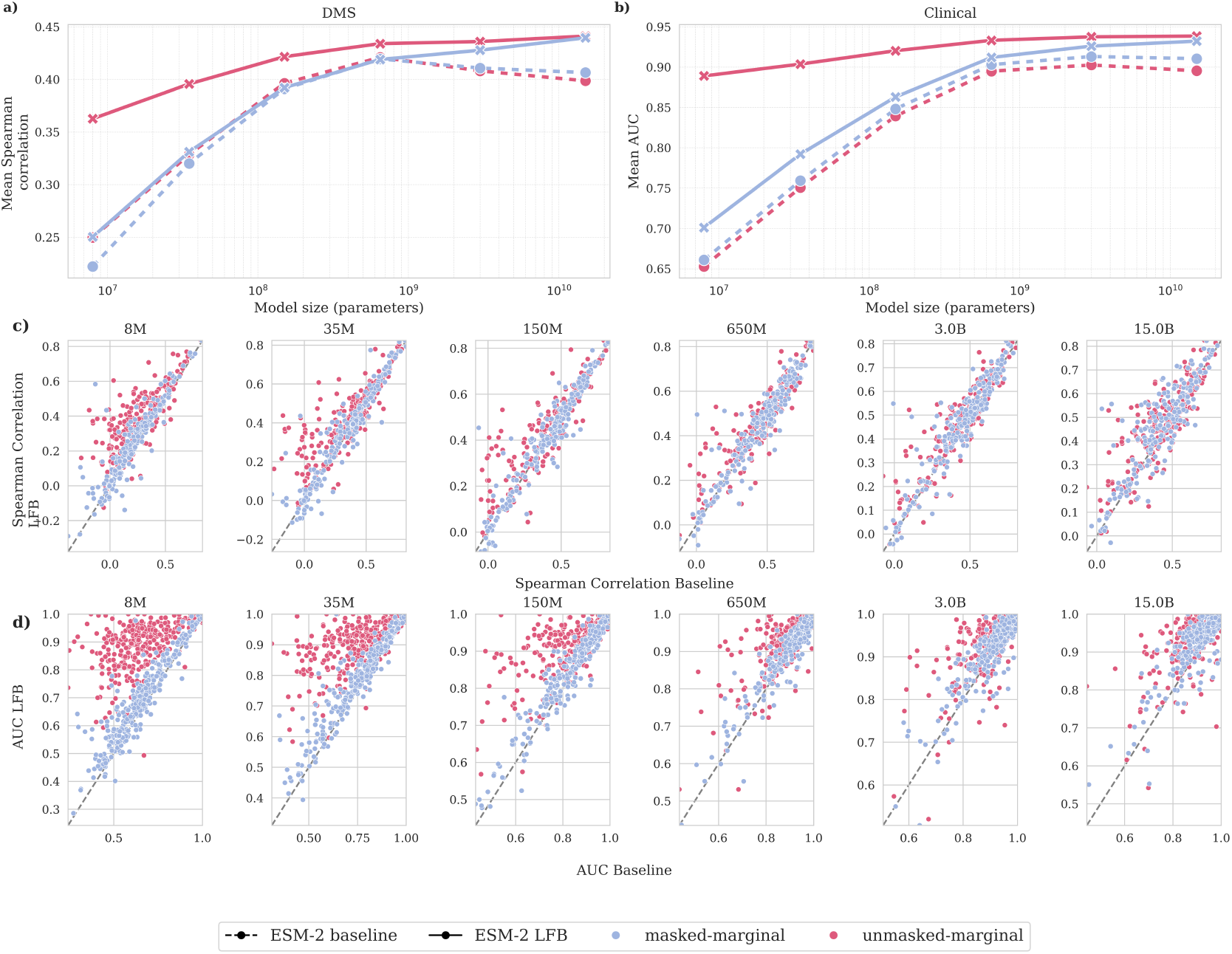
ESM masked-marginal scoring vs unmasked-marginal scoring. a) Average Spearman Correlation to DMS experiments of all models in the ESM-2 family, with masked-marginal scoring versus unmasked-marginal scoring, with and without LFB. b) Average AUC comparison of models in the ESM-2 family, with masked-marginal scoring versus unmasked-marginal scoring, with and without LFB. c) Comparison of all models in the ESM-2 family with masked-marginal scoring versus unmasked-marginal scoring, with and without LFB, on a per-experiment basis. d) AUC performance comparison of of all models in the ESM-2 family with masked-marginal scoring versus unmasked-marginal scoring, with and without LFB, on a per-gene basis. In a) and c), correlations are taken across all the 217 DMS without subsampling of the variants, and the mean is weighted as in Notin et al. [2023].

**Figure F.4.**
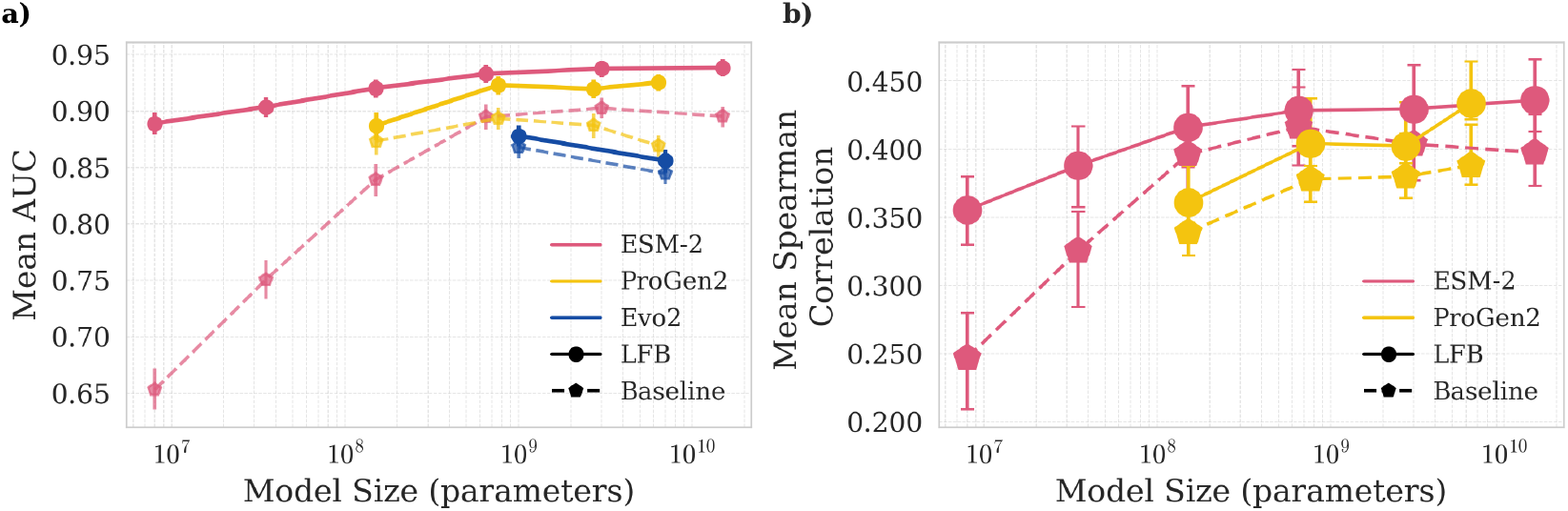
Comparison of pLM and gLM families with and without LFB at clinical label prediction with bootstrap error bars. a) Average AUC comparison of models in ESM-2, ProGen2 and Evo 2 (base) families, with and without LFB. b) Average Spearman Correlation to DMS experiments of all models in ESM-2 and ProGen2 families, with and without LFB, where correlations are taken across all the 217 DMS randomly subsampling to at most 200 variants per assay, and the mean is weighted as in Notin et al. [2023]. Throughout the error bars are bootstrap estimates of the standard deviation of the mean (over roc-auc scores or Spearman correlations) computed by resampling the DMS used or the genes used with replacement.

**Figure F.5.**
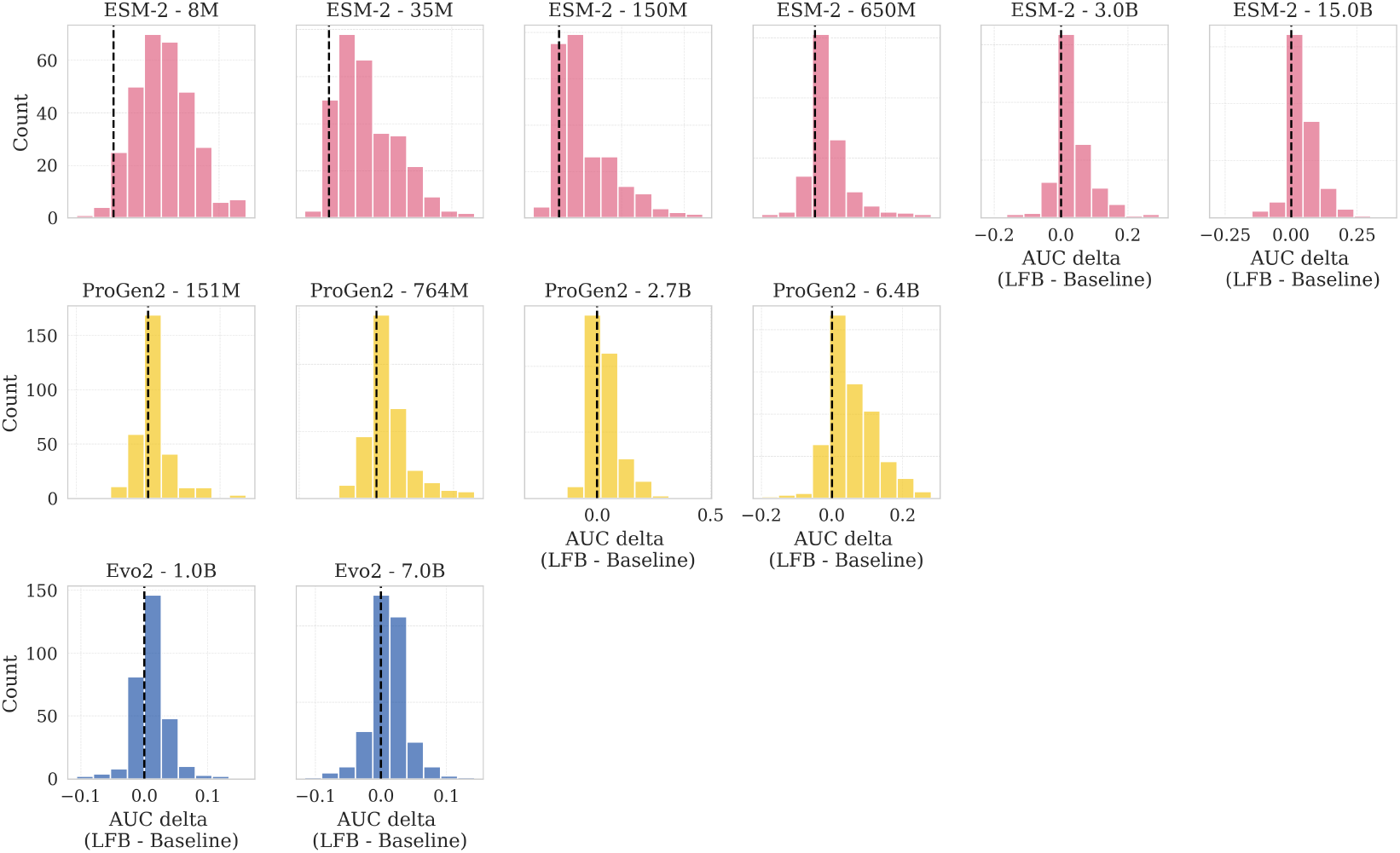
Distribution of performance gains at clinical label prediction using LFB in pLM and gLM, across model families and sizes. Each row corresponds to a model family (ESM-2, ProGen2, Evo 2 (base)), and each column shows models of increasing size (e.g., 8M to 15B parameters). Histograms show the distribution of AUC deltas (Δ AUC = LFB – Baseline) across tested proteins. Vertical dashed lines indicate the zero baseline; bars to the right of the line indicate improved performance with LFB.

**Figure F.6.**
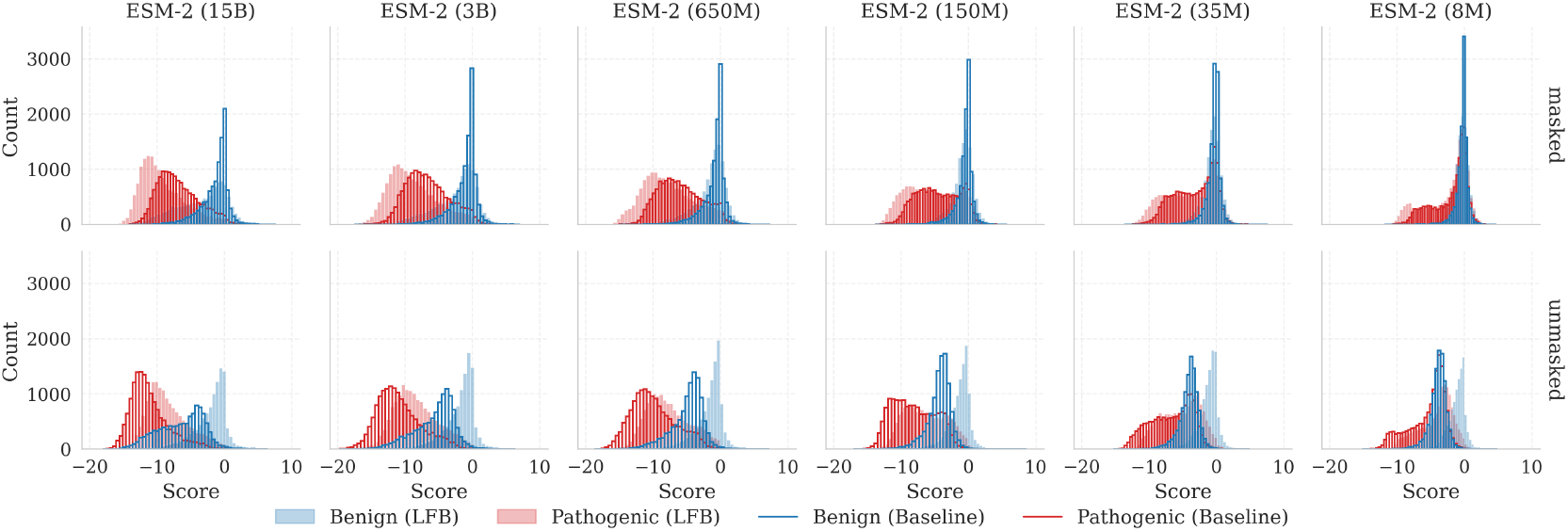
Distributions of scores given to benign and pathogenic labelled variants. Grouping together all (∼ 26, 000) of the benign and pathogenic annotated variants across the 305 genes in the clinical benchmark we show the distributions of scores with (solid) and without (unfilled) LFB for the ESM family of models. The top row shows masked-marginal scoring and the bottom row shows unmasked-marginal scoring.

**Figure F.7.**
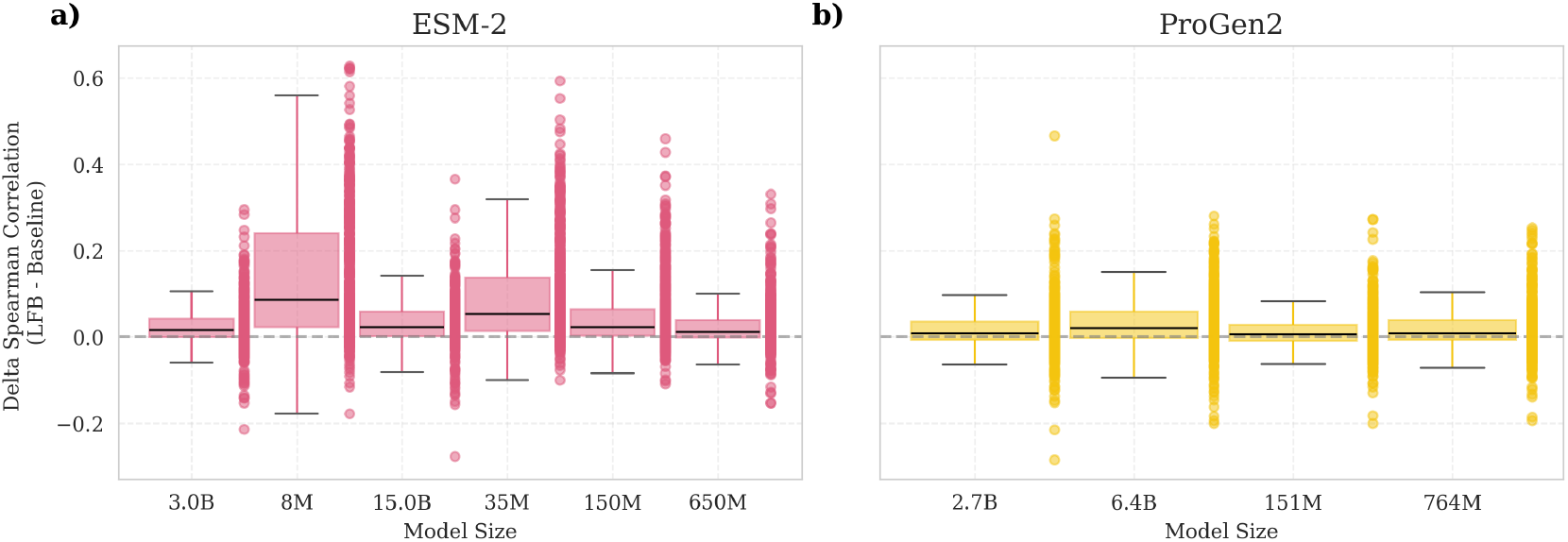
Performance gains at DMS variant prediction using LFB across model sizes for ESM-2 and ProGen2. Each panel displays the difference in Spearman correlation between LFB and baseline predictions across protein deep mutational scanning (DMS) datasets. Boxplots summarize the distribution of deltas for each model size; points represent individual experiments. A horizontal dashed line marks zero difference, with positive values indicating improved agreement with experimental fitness data after applying LFB.

**Figure F.8.**
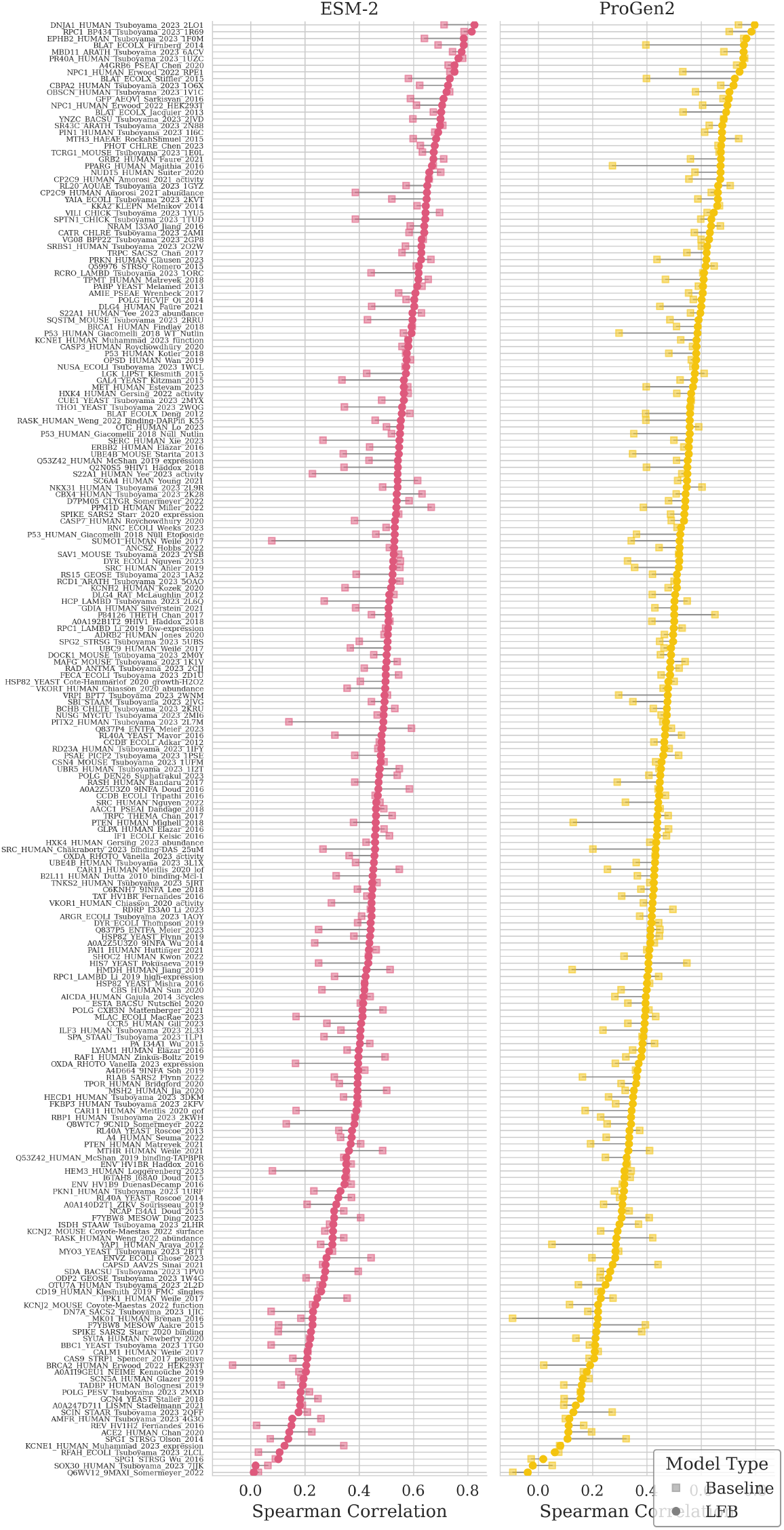
Comparison of models with and without LFB across DMS experiments. Square markers indicate the baseline Spearman correlation, while circular markers represent the LFB-augmented model correlation. Experiments are ordered by increasing LFB correlation within each model. Left panel: ESM-2 (15B), Right panel: ProGen2 (6.4B).

**Figure F.9.**
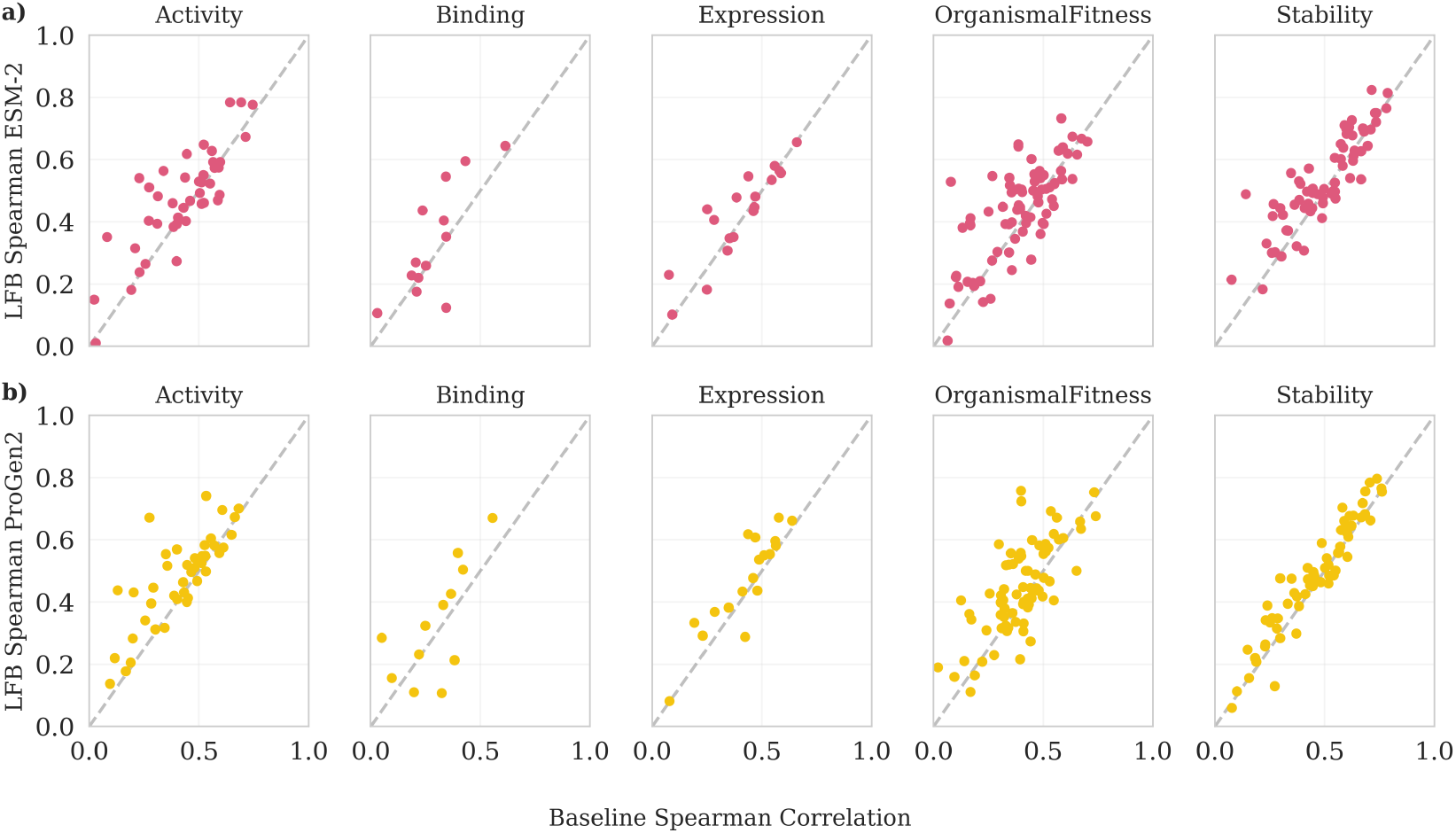
Impact of LFB across distinct DMS functional assays. (a) ESM-2 (15B) LFB, (b) ProGen2 (6.4B) LFB. Each panel represents a different DMS functional assay, grouped by selection type. The x-axis shows the baseline Spearman correlation, while the y-axis represents the LFB-augmented model correlation. The dashed diagonal line indicates the identity line (LFB = Baseline), where no improvement is observed. Points above the diagonal reflect improved correlation with LFB. Across all functional categories, LFB enhances model performance in both ESM-2 and ProGen2 models.

**Figure F.10.**
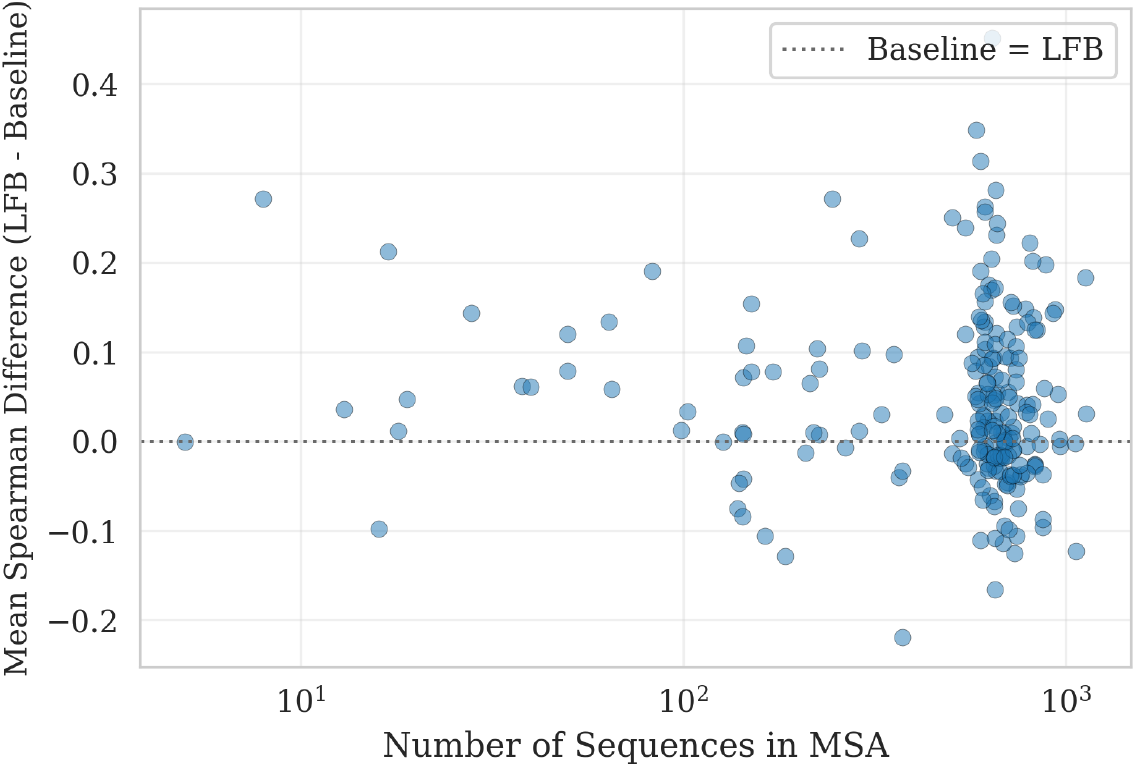
Robustness of likelihood-fitness bridging (LFB) to multiple sequence alignment depth. Relationship between the number of sequences in the multiple sequence alignment (MSA) (log scale, x-axis) and the change in Spearman correlation (LFB - Baseline, y-axis). Each point represents a DMS and the alignment used for LFB averaging produced by MMseqs2 before filtering. The dotted gray line at zero denotes no change between LFB and baseline models, with positive values indicating an improvement in correlation.

**Figure F.11.**
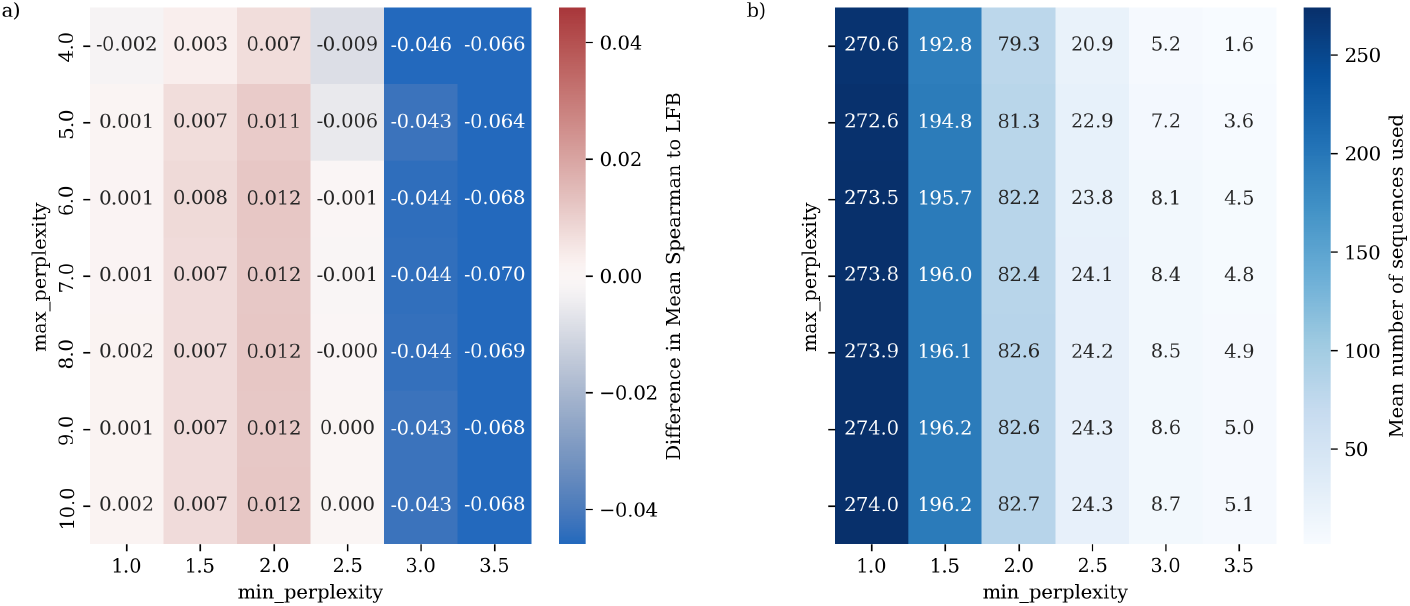
Hyperparameter scan of perplexity based filters on the sequences used for LFB. Comparison between the standard ESM-2 15B LFB model, and LFB estimators obtained by further filtering the sequences used by their minimum pseudo-perplexities and their maximum pseudo-perplexities for a range of thresholds.

**Figure F.12.**
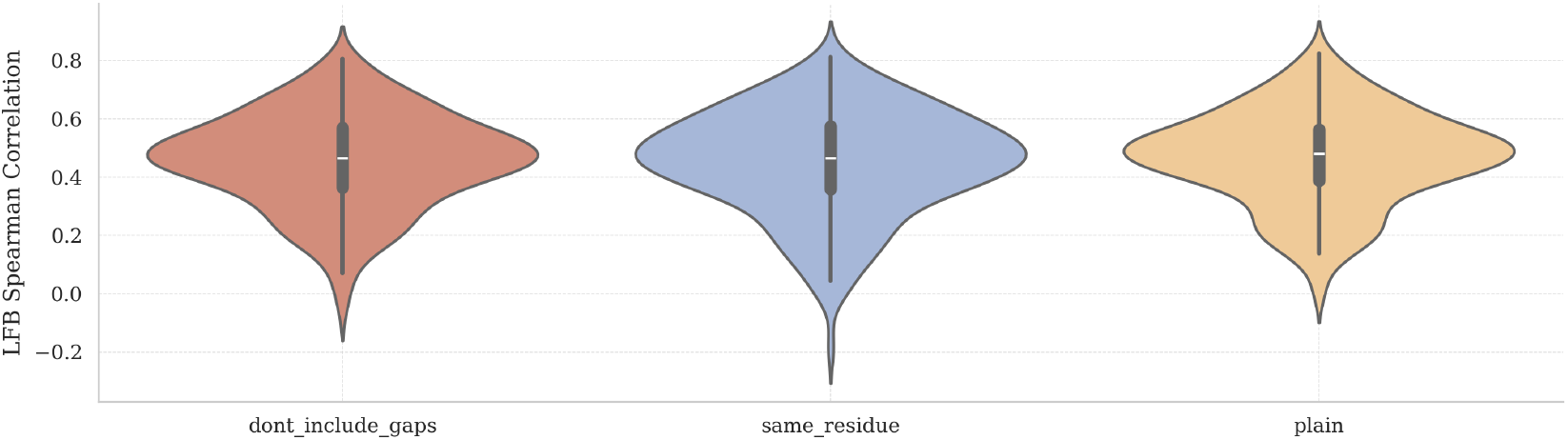
Selected positions for LFB averaging. We show the distributions of Spearman values across DMS assays for three candidate averaging procedures for the ESM-2 15B model: averaging across all sequences for each variant (plain), averaging only across sequences with the same allele as the reference in the variant position (same residue), and averaging only across those sequences which are not a gap position in the variant position (don”t include gaps). Given the slightly higher average, we chose plain to be the standard method for LFB.

## Footnotes

2 Conversely, it has been hypothesized that the likelihood of smaller models can better align with fitness due to a form of model misspecification Weinstein et al. [2022]

3 Throughout we use the same weighting as in [Notin et al., 2023]

## Notes

https://github.com/DiasFrazerGroup/lfb

### Competing Interest Statement

The authors have declared no competing interest.

### Summary of Updates

Updates to section 3.1 on the OU process; Evo 2 40B base model added; slight revision of discussion section; fixed links and typos

